# Ssdp influences Wg expression and embryonic somatic muscle identity in *Drosophila melanogaster*

**DOI:** 10.1101/2021.06.08.447509

**Authors:** Preethi Poovathumkadavil, Jean-Philippe Da Ponte, Krzysztof Jagla

## Abstract

The somatic muscles of the *Drosophila* embryo and larvae share structural and functional similarities with vertebrate skeletal muscles and serve as a powerful model for studying muscle development. Here we show that the evolutionarily conserved Ssdp protein is required for the correct patterning of somatic muscles. Ssdp is part of the conserved Chi/LDB-Ssdp (ChiLS) complex that is a core component of the conserved Wg/Wnt enhanceosome, which responds to Wg signals to regulate gene transcription. *Ssdp* shows isoform specific expression in developing somatic muscles and its loss of function leads to an aberrant somatic muscle pattern due to a deregulated muscle identity program. *Ssdp* mutant embryos fail to maintain adequate expression levels of muscle identity transcription factors and this results in aberrant muscle morphology, innervation, attachment and fusion. We also show that the epidermal expression of Wg is downregulated in *Ssdp* mutants and that Ssdp interacts with Wg to regulate the properties of a subset of ventral muscles. Thus, our data unveil the dual contribution of Ssdp contribution to muscle diversification by regulating the expression of muscle-intrinsic identity genes and by interacting with the extrinsic factor, Wg. The knowledge gained here about Ssdp and its interaction with Wg could be relevant to vertebrate muscle development.

## 1. Introduction

Muscle development is a finely orchestrated process in vertebrates as well as invertebrates involving intrinsic myogenic factors and various signaling molecules that are transduced by downstream effectors into specific gene transcription. An imbalance in any of the proteins involved in this muscle developmental symphony can result in a compensatory mechanism by other players (Mankoo et al. 2003; Relaix et al. 2005; Rudnicki et al. 1993; Kumar et al. 2015) or in case of key factors, might trigger a cascade of deregulation resulting in the disruption of the developmental process (Borello et al. 1999; Lee et Frasch 2000). Vertebrates possess gene families that are often represented by a single orthologue in *Drosophila* (Potthoff et Olson 2007). This correlates with observations of less severe phenotypes in vertebrates when one family member is mutated while mutations in the single *Drosophila* orthologue lead to drastic phenotypes. Patterning by morphogens plays an important role during development. The central dogma of patterning states that morphogen gradients hold positional information that leads to compartmentalization into domains, which then triggers identity acquisition in each domain by the expression of ‘selector’ genes that finally leads to cross-tissue communication and the establishment of new gradients (Lawrence et Struhl 1996). This holds true for vertebrates and invertebrates. In mammals, the evolutionarily conserved morphogen family, Wnt is one of the principal conductors of the developmental processes. It comprises 19 members while *Drosophila* has 7 Wnt homologues including Wingless (Wg). During vertebrate myogenesis, Wnt family members perform non-redundant functions (Münsterberg et al. 1995; Tajbakhsh et al. 1998; Sweetman et al. 2008).

During embryonic skeletal muscle myogenesis in vertebrates, Wnt signaling is among the factors directing the formation of periodically generated somites that form trunk and limb muscles. Soon after formation, ectodermal cues pattern each somite into domains including the high Wnt dermomyotome domain that gives rise to skeletal muscles (Ikeya et Takada 1998; Wagner et al. 2000). Post-mitotic myogenic Pax3^+^ progenitors then delaminate from the dermomyotome to form the myotome by initiating the expression of muscle differentiation genes. In mice, muscle progenitors also express M-Twist (Füchtbauer 1995). These muscle progenitors express specific transcription factors (TFs) depending on their position within the dermomyotome by receiving specific Wnt cues from adjoining tissues. Wnt1 activates MYF5 to form the epaxial myotome that gives rise to back muscles whereas Wnt7a activates MYOD to form the hypaxial myotome that gives rise to muscles of the limb by migrating to limb buds as well as muscles of the ventral body wall, diaphragm and tongue (Tajbakhsh et al. 1998). The initial primary myotome differentiates and elongates anterio-posteriorly in a Wnt11 dependent fashion (Gros, Serralbo, et Marcelle 2009), then undergoes primary myogenesis by fusing with embryonic myoblasts to form primary fibers by acquiring a specific muscle identity that determines its morphology, innervation, attachment and function. The factors that define the identity of individual muscles that distinguish them from their neighbors are yet to be determined in vertebrates.

*Drosophila* somatic or body wall muscles are similar to vertebrate skeletal muscles since they are syncytial, striated and voluntary. *Drosophila* has 30 larval somatic muscles per hemisegment arranged in a stereotypical pattern generated during embryonic myogenesis. *Drosophila* embryos undergo simultaneous and synchronous segmentation of the germ layers giving rise to parasegments in a Wingless (Wg) dependent fashion (Bejsovec et Martinez Arias 1991). They are subsequently divided into domains including the high Wg (epidermal cue), high Twist (Twi) mesodermal domain that generates larval somatic muscles (Lee et Frasch 2000). Muscle progenitors are subsequently generated that then divide asymmetrically to give rise to founder cells (FCs), each FC containing the information for one specific muscle. Each FC expresses a muscle identity transcription factor (iTF) code that dictates its identity. Ectodermal Wg cues are implicated in the specification of some Slou^+^ ventral muscle progenitors (Cox, Beckett, et Baylies 2005). It is not known if Wnt signals play a role in regulating gene expression in specific muscles at later stages.

Although a broad requirement for Wnt signaling at various stages of muscle development has been identified, various factors involved in transducing this signal and translating it into gene transcription at each timepoint as well as those regulating its own expression remain elusive. *Sequence-specific single-stranded DNA-binding protein* (*Ssdp*) is an evolutionarily conserved gene with homologues across the animal kingdom (Bayarsaihan, Soto, et Lukens 1998; Castro et al. 2002; Chen et al. 2002; van Meyel, Thomas, et Agulnick 2003). It is part of the conserved Chip/LDB-Ssdp (ChiLS) complex along with the transcriptional cofactor known as Chip (Chi) in *Drosophila* and LIM domain-binding protein 1 (LDB1) in humans (H. Wang et al. 2019, 2020). LDB1 is required for the nuclear localization of Ssdp (van Meyel, Thomas, et Agulnick 2003) and binds LIM homeodomain TFs with high affinity while Ssdp protects LDB1 from proteasomal degradation (Güngör et al. 2007; Y. Wang et al. 2010; Bronstein et al. 2010). ChiLS has recently been shown to constitute a core component of the Wnt enhanceosome that translates the conserved canonical Wnt signaling mediated by ß-catenin into gene transcription (Fiedler et al. 2015; Renko et al. 2019). Humans possess 3 *Ssdp* homologues (*SSBP2, SSBP3* and *SSBP4*) while *Drosophila* has only one. This gene family is of interest for the role of its members in human cancers since they are mis-regulated in multiple forms of cancer (Liu et al. 2008; Poitras et al. 2008; Y. Wang et al. 2010), as is the case for Wnt signaling components (Delgado-Deida, Alula, et Theiss 2020).

The role of Ssdp during myogenesis has not yet been investigated, neither have its spatial and temporal roles. Here, we identify *Ssdp* as a significantly differentially regulated gene using a muscle-subset-specific Translating Ribosome Affinity Purification (TRAP) approach and aim to dissect the consequences of its loss of function on *Drosophila* embryonic muscle development. We show that *Ssdp* expression is enriched in somatic muscles during mid to late stages of muscle development. Its loss of function affects the levels of expression of muscle iTFs and influences the acquisition of muscle identity with most severe defects in the ventral and lateral muscles. Considering the known role of Ssdp in Wg-dependent gene regulation and the partial overlap of *Ssdp* and *wg* loss of function muscle phenotypes, we propose that in addition to its intrinsic role in maintaining iTF expression, Ssdp also ensures late Wg function in muscles.

## 2. Materials and Methods

### Drosophila strains

All stocks except the temperature sensitive *wg^ts^* were grown at a temperature of 25°C. The following stocks were gifted to us: *Ssdp^L5^* and *Ssdp^L7^* (gifts from Donald J van Meyel, McGill Centre for Research in Neuroscience, Montreal, Canada (van Meyel, Thomas, et Agulnick 2003)), S59-Gal4 (gift from Manfred Frasch, Erlangen, Germany). The following strains were obtained from Bloomington *Drosophila* Stock Center: *wg^ts^* is a heat sensitive amorphic allele (wg[I-12]bw[1]/Cyo) that was rebalanced on a CyO(Act-GFP) balancer to distinguish homozygotes,

24B-Gal4 (w[*]; P{w[+mW.hs]=GawB}how[24B]),
w[1118]; PBac{w[+mC]=IT.GAL4}Ssdp[2082-G4]/TM6B Tb[1],
lms-Gal4 (w[1118]; P{y[+t7.7] w[+mC]=GMR88F08-GAL4}attP2),
UAS-LAGFP (y[1] w[*]; P{y[+t*] w[+mC]=UAS-Lifeact-GFP}VIE-260B) and
UAS-RpL10a-GFP (w[*]; P{w[+mC]=UAS-GFP-RpL10Ab}BF2b).

The following stock was obtained from Kyoto Stock Center: w[*]; P{UAS-Act5C.T:GFP}10-2

We used the following genotypes: 24B>Gal4;UAS>dTCF^DN^, S59>Gal4/MKRS;UAS>RpL10a-GFP and lms>Gal4; UAS>LAGFP.

### Temperature shift experiments

Temperature shift experiments were conducted as follows: To determine the role of Wg during early stages of muscle development, *wg^ts^* flies were grown at a permissive temperature of 18°C for 12 hours (mid stage 11) after which the apple juice plates with embryos were shifted to a non-permissive temperature of 28°C for 9 more hours before fixing them. To determine the role of Wg at slightly later stages, embryos were staged by letting flies lay eggs for 3 hours at 18°C after which the apple juice plates were collected and let to develop at 18°C for 14 more hours and subsequently shifted to 28°C for 5 hours.

### Immunofluorescent staining

The following primary antibodies were used: rat anti-Actin (1/500, MAC 237; Babraham Bioscience Technologies), rabbit anti-β3 Tubulin (1:5000; R. Renkawitz-Pohl, Philipps University, Marburg, Germany), rat anti-Tropomyosin (1:200, ab50567, Abcam), mouse anti-FasII (1:500, 1D4, Developmental Studies Hybridoma Bank (DHSB)), anti-GFP (1:1000, DHSB), mouse anti-βPS integrin (1:200 DSHB), mouse anti-Col (1:50, from Alain Vincent, Center for Integrative Biology, Toulouse, France), rabbit anti-Mef2 (1:500, from Eileen Furlong, EMBL, Germany), rabbit anti-Slou (1:300, from Manfred Frasch, Erlangen, Germany) and mouse anti-Wg (1:500, 4D4, DHSB). Fluorescence conjugated secondary antibodies from Jackson ImmunoResearch produced in donkey conjugated with Alexa 488, Cy3 or Cy5 were used at a concentration of 1:300. For immunostainings using primary antibodies produced in rat and mouse, minimal cross secondary antibodies against both species were used.

### RNA FISH and in situ hybridization

For RNA FISH experiments (Raj et al. 2008), 29 Quasar 570 conjugated Stellaris probes from LGC Biosearch Technologies (Orjalo, Johansson, et Ruth 2011) targeting the long isoforms of Ssdp were used. Fixed embryos were used to hybridize the probes using the standard Stellaris hybridization procedure. This was followed by antibody staining against actin to visualize all muscles and GFP to visualize muscle subsets.

For *in situ* hybridization, we used the following primer pairs:

*Ssdp*: 5’-TGTACGAATATCTGCTGCACG-3’ and
5’-GCATCGTCGAGTTAGGGAAG-3’

Probe generation and *in situ* hybridization were performed following the standard procedure (Legendre et al. 2013). Reverse primers with a T7 promoter sequence prefix were used. PCR fragments were amplified using the genomic DNA as template, purified and reverse transcribed using the Roche SP6/T7 transcription kit with Dig-UTP to generate Digoxygenin labelled mRNA probes. Fixed embryos were then hybridized *in situ* with the probes followed by TSA amplification and antibody staining against actin.

### Image acquisition, processing and statistical analysis of images

All images were acquired on a Leica SP8 microscope using a 40X objective at a resolution of 1024×1024 or 2048×2048. They were analyzed and processed using ImageJ. Statistical tests and graph generation were performed in R. For CTCF quantification of fluorescence intensities of Col and Slou in WT versus *Ssdp* mutants, 25 stacks were acquired for each embryo analyzed. An equal number of images at 1024×1024 and 2048×2048 were included in each group to be compared against. ROIs were manually selected on maximum projections of each image and the mean of three areas close to each ROI was used as the background fluorescence. To quantify the number of Mef2+ nuclei, nuclei were manually counted in each DT1 muscle in abdominal hemisegments A2-A5 for each embryo. Similarly, the number of Eve+ pericardial cells were manually counted in hemisegments A2-A5 of each embryo analyzed.

### Statistical analysis of transcriptomic data and cis regulatory motif analysis

Differential gene expression of the transcriptomic microarray data was determined in R using the limma package. Enrichment for cis regulatory motifs (CRMs) was determined using i-cisTarget (Herrmann et al. 2012; Imrichová et al. 2015). A normalized enrichment score (NES) threshold of 3 was used. Graphs were generated using R.

## 3. Results

### 3.1. Ssdp mRNA under translation is differentially expressed in muscle subsets

We used TRAP data generated earlier for two somatic muscle subsets expressing distinct iTFs, one expressing Slouch (Slou/S59) (S. Knirr, Azpiazu, et Frasch 1999) and the other expressing Lateral muscles scarcer (Lms) (Müller et al. 2010; Bertin et al. 2015, 2021) as well for the global muscle population expressing Duf. We also generated transcriptomic data for the entire embryo. Data analysis revealed that *Ssdp* was among the genes that were significantly upregulated in the Lms subset during late stages (13-16 hours after egg laying or AEL that we refer to as time window 3 or T3 here) (Figure 1A). We also observed a significant enrichment for CT-rich as well as complementary GA-rich cis regulatory motifs among the genes significantly upregulated during this time window (Figure 1B). Chicken SSDP was initially identified as a protein capable of binding CT-rich tracts in the α2(I) collagen gene. These tracts were subsequently shown to be capable of binding the fly Ssdp protein (Bayarsaihan, Soto, et Lukens 1998; Bronstein et al. 2010). The expression of *Ssdp* mRNA under translation mapping to all transcripts showed an upward trend in both muscle subsets with respect to the global embryonic mRNA. When compared with translating mRNA in the global muscle population, it is significantly upregulated in the Lms subset with respect to the Slou subset (Figure 1C-E).

**Figure 1.**
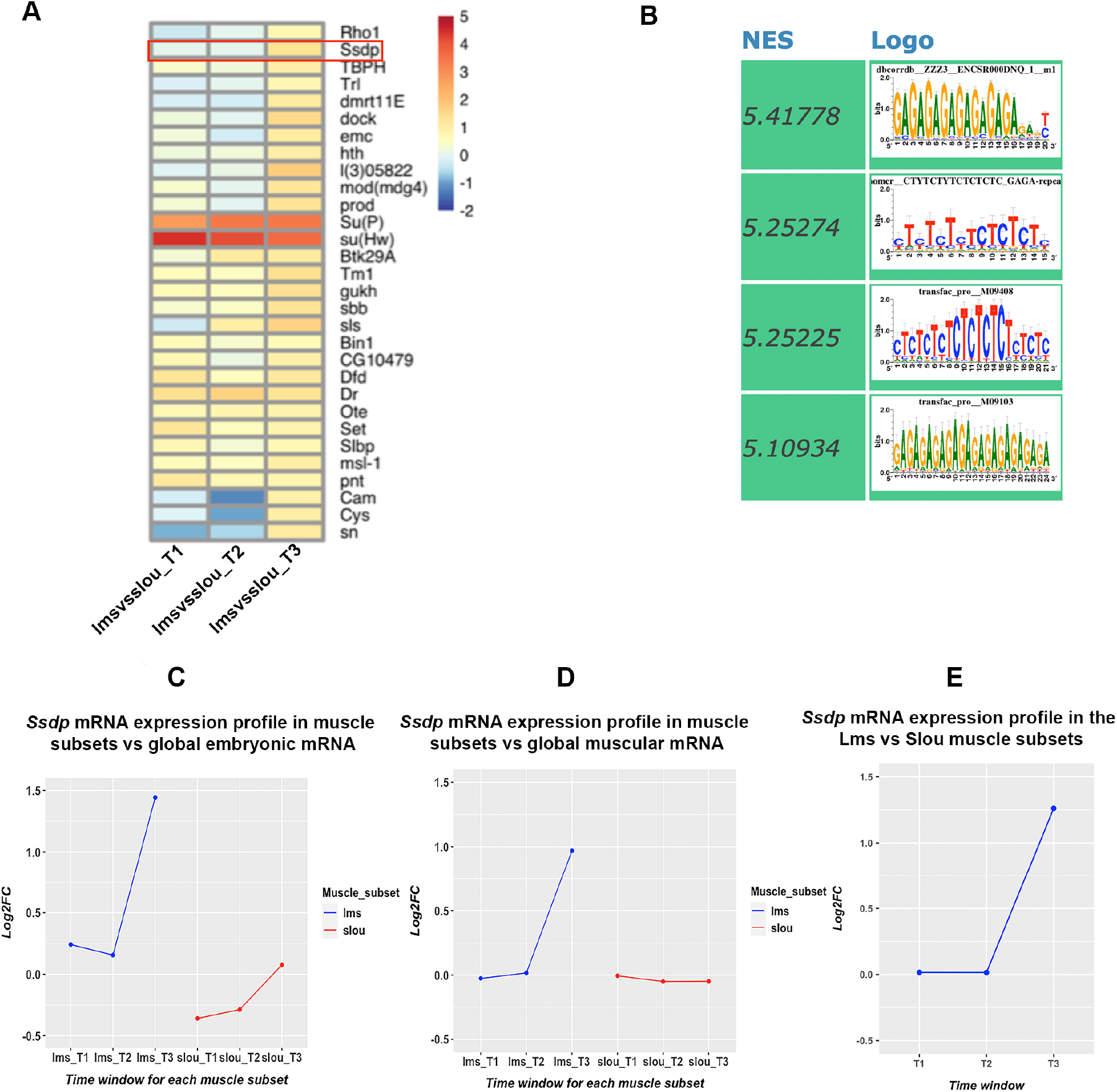
*Ssdp* mRNA undergoing translation display significant differential expression in muscle subsets at late embryonic stages. **(A)** *Ssdp* is among the genes that are significantly upregulated in the Lms muscle subset during time window T3 that represents late embryonic stages. **(B)** An i-cisTarget analysis of cis regulatory regions among genes significantly upregulated in the Lms subset versus the Slou subset during T3 reveals a significant enrichment for CT-rich and complementary GA-rich motifs (*NES = Normalized Enrichment Score calculated by i-cisTarget above a threshold value of 3*). **(C-E)** The expression profile of *Ssdp* mRNA undergoing translation derived from TRAP data shows that it displays an upward trend in the Lms as well as Slou muscle subsets with respect to the global embryonic mRNA (C) and is significantly upregulated in the Lms population with respect to the global Duf+ muscle population (D). This is confirmed by a comparison of the muscle subsets amongst themselves (E).

Since Ssdp is an evolutionarily conserved protein, we examined its alignment with the canonical isoforms of human SSBP proteins. Apart from the already recognized conservation in the LUFS domain, we observed three other smaller blocks with highly conserved amino acid residues (Supplementary figure 1). One of the conserved regions is part of a proline-rich region whose deletion led to a headless phenotype in mice (Enkhmandakh, Makeyev, et Bayarsaihan 2006) due to the loss of the fore and midbrain. No function, if any, has been attributed to the other conserved regions yet.

### 3.2. Ssdp mutants display severe somatic muscle phenotypes

Given the differential expression observed in our TRAP datasets, we were interested to see if this gene played specific roles in individual muscles during muscle diversification. To this end, we analyzed *Ssdp^L5^* and *Ssdp^L7^*, both considered to be *Ssdp* null mutants. *Ssdp^L7^* is a deletion of exon 2 that contains the complete Ssdp protein coding sequence while *Ssdp^L5^* is a partial deletion of this exon (van Meyel, Thomas, et Agulnick 2003). We observe that both mutants display similar severe muscle phenotypes as well as concomitant muscle innervation defects (Figure 2).

**Figure 2.**
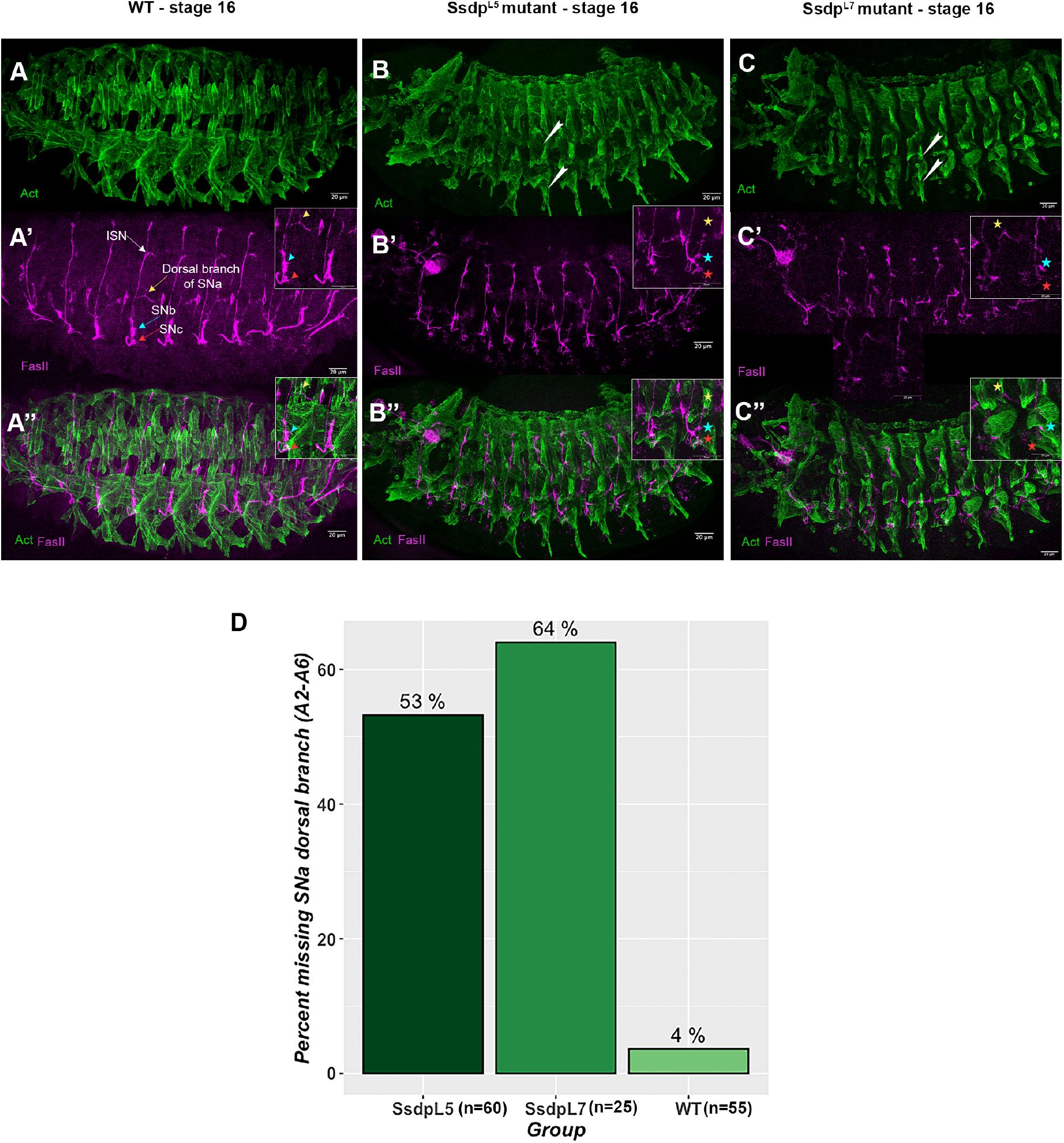
Loss of function of *Ssdp* leads to severe defects in the somatic muscles. **(A-A”)** The muscle (A) and innervation (A’) pattern in WT stage 16 embryos. Different motor neurons are indicated by different colored arrows. Insets in A’ and A” show the stereotypical innervation and muscle patterns in each hemisegment. **(B-B”)** Both the muscle pattern (arrowheads in B) and innervation (asterisks in the inset in B’) are severely defective in *Ssdp^L5^* mutants, which presumably have a partial deletion in the *Ssdp* gene that contains a single protein coding exon. The lateral and ventral muscles are severely affected as indicated by arrowheads in B. The insets in B’ and B” display zoomed views highlighting innervation defects such as morphologically defective SNb and SNc motor neurons and a missing dorsal branch of the SNa. Compare asterisks to similarly colored arrowheads in the WT inset in A’. **(C-C”)** *Ssdp^L7^* mutants lacking the entire length of the *Ssdp* gene display phenotypes resembling *Ssdp^L5^* mutants with muscle morphology (arrowheads in C) and innervation (asterisks in the inset in C’, C”) defects. **(D)** The percentage of hemisegments (A2-A6) missing the dorsal branch of the SNa is slightly higher in *Ssdp^L7^* mutants compared to *Ssdp^L5^* mutants, although it is significantly higher than in WT embryos in both mutants.

In the late stage WT embryo, somatic muscles are arranged in a stereotypical pattern. In both *Ssdp^L5^* and *Ssdp^L7^* mutants, however, although muscles are generally present, individual muscle fibers appear compacted and/or of aberrant morphology (Figure 2A, B, C). This is particularly obvious in lateral and ventral regions. Aberrations in the innervation of lateral and ventral muscles by their specific motor neurons are also apparent. The dorsal branch of the SNa motor neuron that innervates the WT lateral transverse (LT) muscles is missing in some segments in stage 16 *Ssdp^L5^* and *Ssdp^L7^* mutants (Figure 2A’, B’, C’). Similarly, the SNb and SNc branches normally innervating ventral muscles are severely affected with irregular morphologies and non-uniform innervation patterns in different hemisegments while the ISN branch targeting dorsal muscles shows only minor trajectory defects. The percentage of hemisegments where the dorsal branch of the SNa fails to defasciculate is slightly more pronounced in *Ssdp^L7^* mutants compared to *Ssdp^L5^* mutants (Figure 2D).

These observations reveal that Ssdp is required for proper patterning and innervation of somatic muscles with a major impact on ventral and lateral muscles. Our subsequent analyses were performed on *Ssdp^L5^* flies since both mutants display similar phenotypes and this line is easier to amplify. Henceforth, we will refer to the homozygous *Ssdp^L5^* embryos as ‘*Ssdp* mutants’ and explicitly refer to *Ssdp^L7^* where applicable.

### 3.3. Ssdp mRNA is expressed in somatic muscles

In order to analyze embryonic expression patterns of *Ssdp*, we used *in situ* hybridization to reveal its transcripts and a transgenic *Ssdp* enhancer trap line in which GAL4 expression is driven by *Ssdp* regulatory sequences, thus giving an indication of the tissues in which Ssdp is expressed and its expression levels. We first performed RNA FISH (Raj et al. 2008) with probes that map to the 3’ UTR of the two long *Ssdp* isoforms (Figure 3). *Ssdp* transcripts for the long isoforms cannot be detected in the somatic mesoderm during the specification of muscle precursors/founders (stage 11-12). However, later in development (stage 15), *Ssdp* mRNAs accumulate in all somatic muscles. In stage 15 and early stage 16 embryos, *Ssdp* transcripts are uniformly distributed in muscles as well as the ventral nerve chord (VNC) (Figure 3B-D’). Thus, using RNA FISH, we do not detect a particular enrichment of *Ssdp* transcripts in any muscle subset. However, this analysis represents whole muscular mRNA as opposed to the TRAP transcriptomic data that aimed to discover mRNA under translation. Also, the RNA FISH probes map to the two long *Ssdp* isoforms with long 3’UTR regions, and could thus present only a partial picture of the *Ssdp* expression pattern.

**Figure 3.**
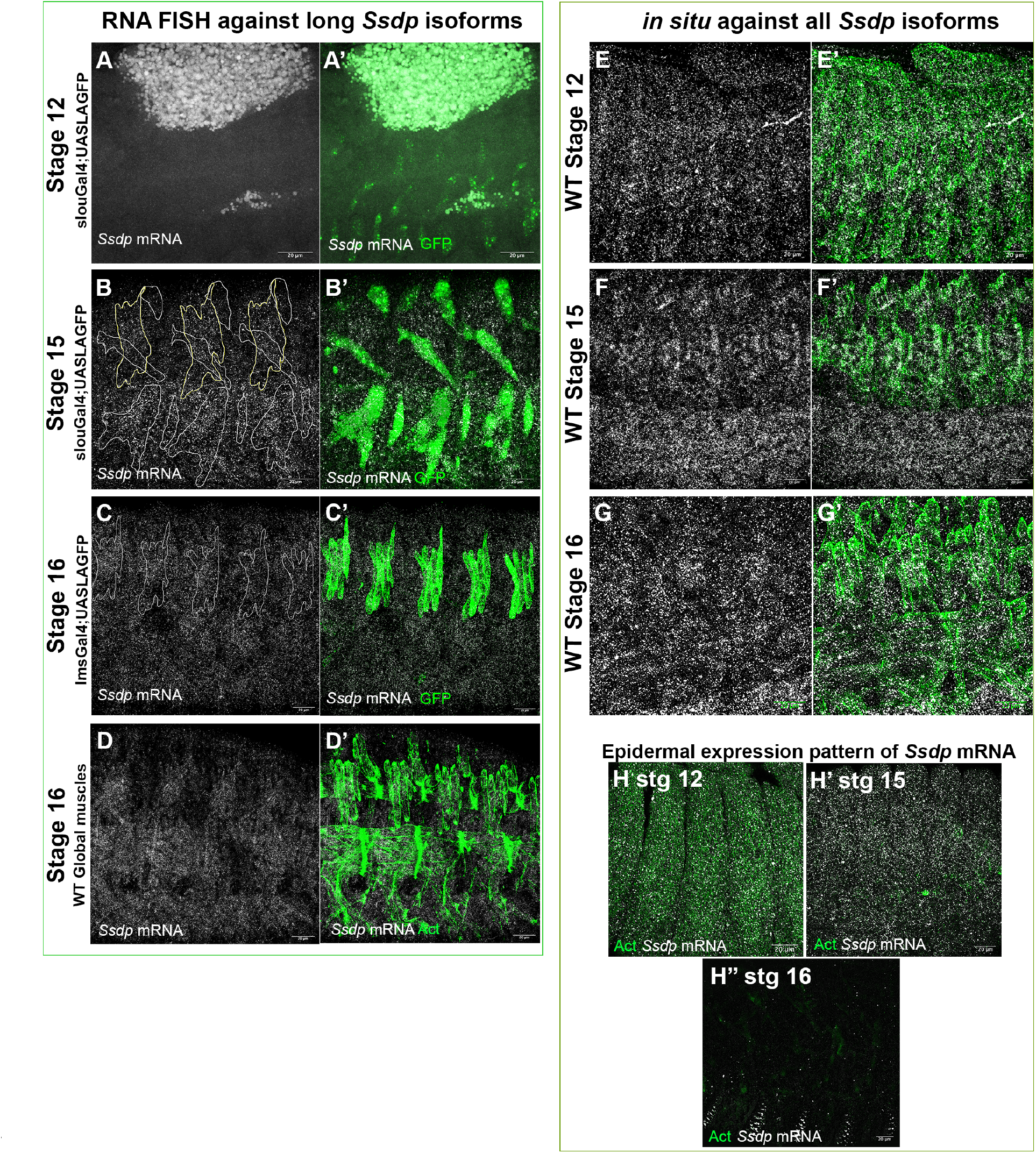
RNA FISH against the long *Ssdp* isoforms reveals muscular expression at late stages whereas an *in situ* hybridization targeting all isoforms also reveals expression at early and very late stages. **(A-A’)** No remarkable transcript expression is detected during stage 12. The Slou+ muscle subset is visualized by an anti-GFP antibody as revealed by the Slou-Gal4 driven expression of RpL10aGFP. (C’) **(B-B’)** Stage 15 embryos show high muscular expression levels of *Ssdp* mRNA. The Slou and Lms muscle subsets are outlined in white and yellow respectively in (B) as revealed by an antibody against actin (not shown). **(C-C’)** *Ssdp* expression persists in stage 16 embryos. The Lms+ lateral transverse (LT) muscles are outlined in (C) as revealed by LifeActGFP driven by the lms-Gal4 driver. **(D-D’)** *Ssdp* expression in global somatic muscles in stage 16 embryos as revealed by an antibody against actin. **(E-E’)** *in situ* hybridization targeting all *Ssdp* isoforms reveals mesodermal expression at stage 12 that is absent for long isoforms. **(F-G’)** The somatic muscles and ventral nerve chord (VNC) express high levels of *Ssdp* at stage 15 (F-F’) and stage 16 (G-G’). (H-H”) *in situ* reveals epidermal expression that is not present for long isoforms. While this expression is throughout the epidermis during stages 12-16, by very late stage 16, expression becomes restricted to a characteristic segmentally repeated pattern of anterio-ventral epidermal stripes.

To determine the expression pattern of all isoforms, we performed a classic *in situ* hybridization with a *Ssdp* probe targeting a region of the protein coding exon of *Ssdp* present in all transcripts. In addition to the pattern described above with *Ssdp* expression in the VNC and developing muscles, we also detect *Ssdp* transcripts in epidermal and mesodermal layers at stage 12 (Figure 3E-H”). This also reveals epidermal expression during early to late stages followed by a segmentally repeated expression in ventral epidermal stripes during late stage 16 that is not observed for the long isoforms. Thus, in contrast to the TRAP profiles, neither the probes targeting long *Ssdp* isoforms nor those targeting all its isoforms detect an enrichment in particular muscle subsets. This suggests that the TRAP dataset could reveal a muscle-subset specific regulation of *Ssdp* at the level of mRNA undergoing translation. Due to the unavailability of a Ssdp antibody, we are unable to confirm this possibility. A Ssdp-Gal4 driven expression of GFP tagged actin5C reveals a muscular expression pattern that increases over time with marked GFP accumulation at stage 15 in Lms-positive LT muscles and Slou-positive DT1 (Supplementary Figure 2). This developmentally regulated expression suggests stage and isoform specific roles for Ssdp in somatic muscles.

### 3.4. Cytoskeletal muscle components are disorganized in the absence of Ssdp

Actin is a key muscle protein that is an integral component of sarcomeres, the contractile units of the muscle, apart from playing its generic role in the actin cytoskeleton (A. F. Huxley et Niedergerke 1954; H. Huxley et Hanson 1954). Actin dynamics during developmental stages 12-15 are involved in the formation of the fusogenic synapse that permits fusion of fusion competent myoblasts (FCMs) with the myotube (Sens et al. 2010; J. H. Kim et al. 2015). During muscle attachment, it plays a crucial role in extending filopodia to sense correct attachment sites (Schnorrer et Dickson 2004; Richier et al. 2018). During muscle innervation, muscles extend myopodia to communicate and connect with the right presynaptic filopodia (Ritzenthaler, Suzuki, et Chiba 2000). Thus, any disruptions to actin and its cofactors are potentially detrimental to muscle development. We examined the expression pattern of actin as well as its binding partner and muscle differentiation marker Tropomyosin (Tm). Both proteins are expressed in *Ssdp* mutants. Actin presents a highly organized arrangement in WT stage 16 embryos. Notably, it is cortically enriched outlining lateral transverse (LT) muscle shapes. This actin distribution is lost in *Ssdp* mutants in which actin-stained individual LT muscles are difficult to distinguish (Figure 4A, B). Similarly, the muscle differentiation marker Tm2 that is implicated in myotube elongation by co-localizing with F-actin (Williams et al. 2015) and enriched in LT muscle termini in WT embryos, displays a fuzzy, irregular pattern in *Ssdp* mutant LTs that fail to fully elongate (Figure 4C, D).

**Figure 4.**
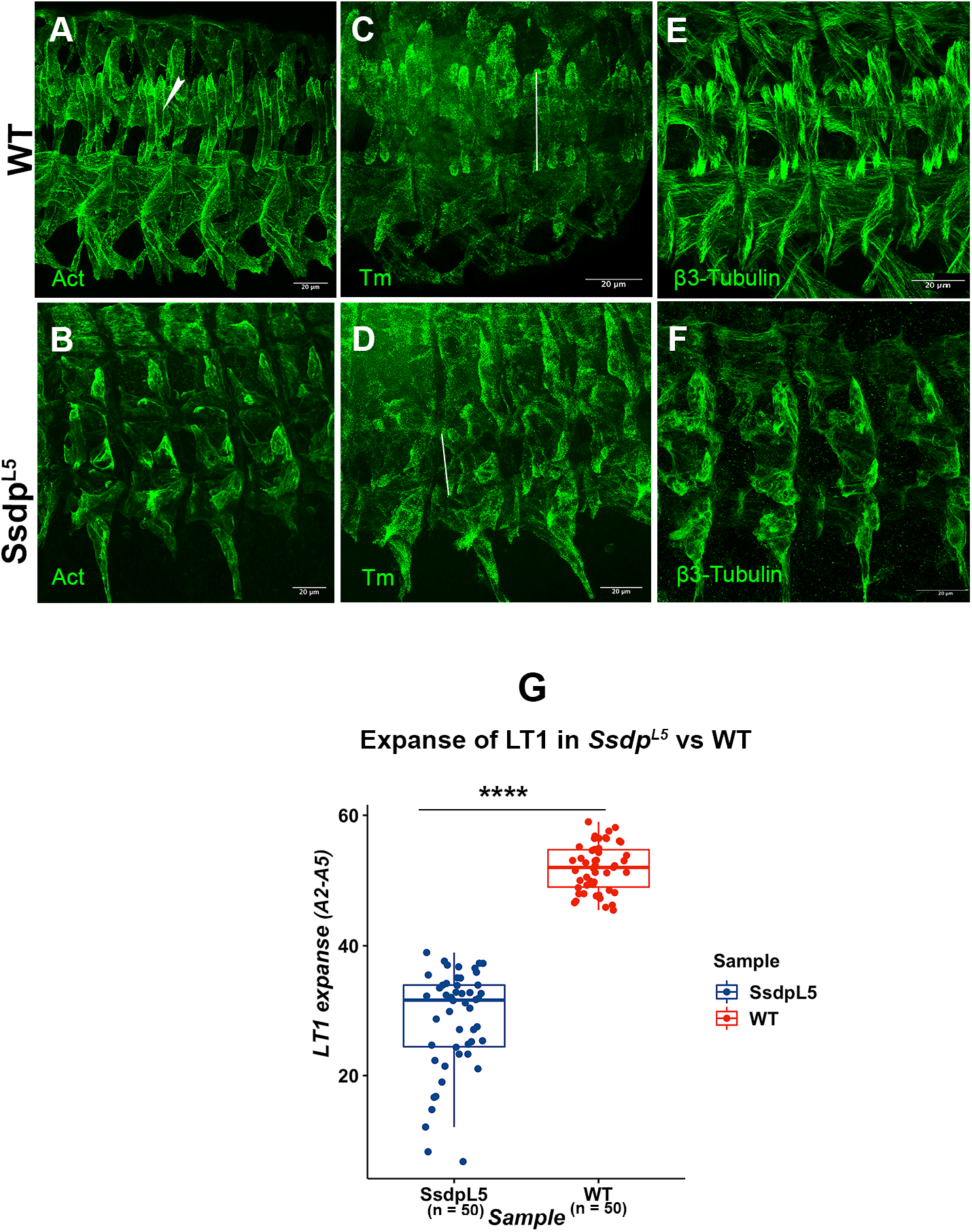
*Ssdp* mutants exhibit severe disorganization of the actin cytoskeleton and microtubules at stage 16. **(A-B)** In WT embryos (A), a staining for actin using an anti-actin antibody reveals a structured organization of the actin cytoskeleton whereas this organization is severely disrupted in *Ssdp* mutants with a disorganized concentration of actin at muscle ends (B). **(C-D)** A disorganization similar to the actin network is observed for Tm2, an actin binding protein and muscle differentiation marker, where WT embryos have an organized arrangement (C) as opposed to *Ssdp* mutants (D). **(E-F)** The microtubule network as visualized by an antibody against ß3-Tubulin shows that this network is equally disorganized in *Ssdp* mutants (F) in comparison to WT embryos (E). **(G)** The expanse of the distance to which the LT1 muscles extend in each hemisegment is significantly lower in *Ssdp* mutants as indicated by a t-test. **** = p-value < 0.0001 at a 95% confidence interval. The expanse measured for each LT1 is indicated by a line in (C) and (D).

Another essential cytoskeletal muscle component is the microtubule (MT) network. MT associated proteins such as dynein play a crucial role in determining muscle length and myonuclear positioning in the LT muscles (Folker, Schulman, et Baylies 2012). WT LT muscles extend longitudinally in both directions with ß3-Tubulin accumulating at LT extremities. In *Ssdp* mutants, ß3-Tubulin enrichment at LT ends cannot be detected and LT muscles extend over much shorter distances compared to the WT with muscle ends bending towards each other (Figure 4E, F). We observe that the LT1 muscle extends over around half the distance compared to the WT (Figure 4G).

Since the muscle differentiation marker, Tm2 is expressed in somatic muscles and muscles are arranged more or less in their WT pattern although they are severely affected, this indicates that the muscles initiate the differentiation program, but fail to establish their identity and the identity program is possibly deregulated.

### 3.5. iTF expression is downregulated in the absence of Ssdp

Muscle identity acquisition is regulated by iTFs and their downstream realisators. We previously demonstrated that the attenuated expression of one iTF, Ladybird (Lb) (Junion et al. 2007) causes perturbations in the identity acquisition of the Lb-dependent segment border muscle (SBM). The affected muscle pattern and innervation observed in *Ssdp* mutants prompted us to test whether the loss of *Ssdp* could have an impact on the expression of iTFs and in turn the acquisition of muscle identity. We chose to test two iTFs, Collier (Col) determining dorsal DA3 muscle identity (Crozatier et Vincent 1999) and Slou involved in the identity of several ventral and lateral muscles including the ventral acute VA2/3, dorso-lateral DT1 and ventral transverse VT1 muscles. The expression of Col as well as Slou is attenuated in a *Ssdp* loss of function context (Figure 5).

**Figure 5.**
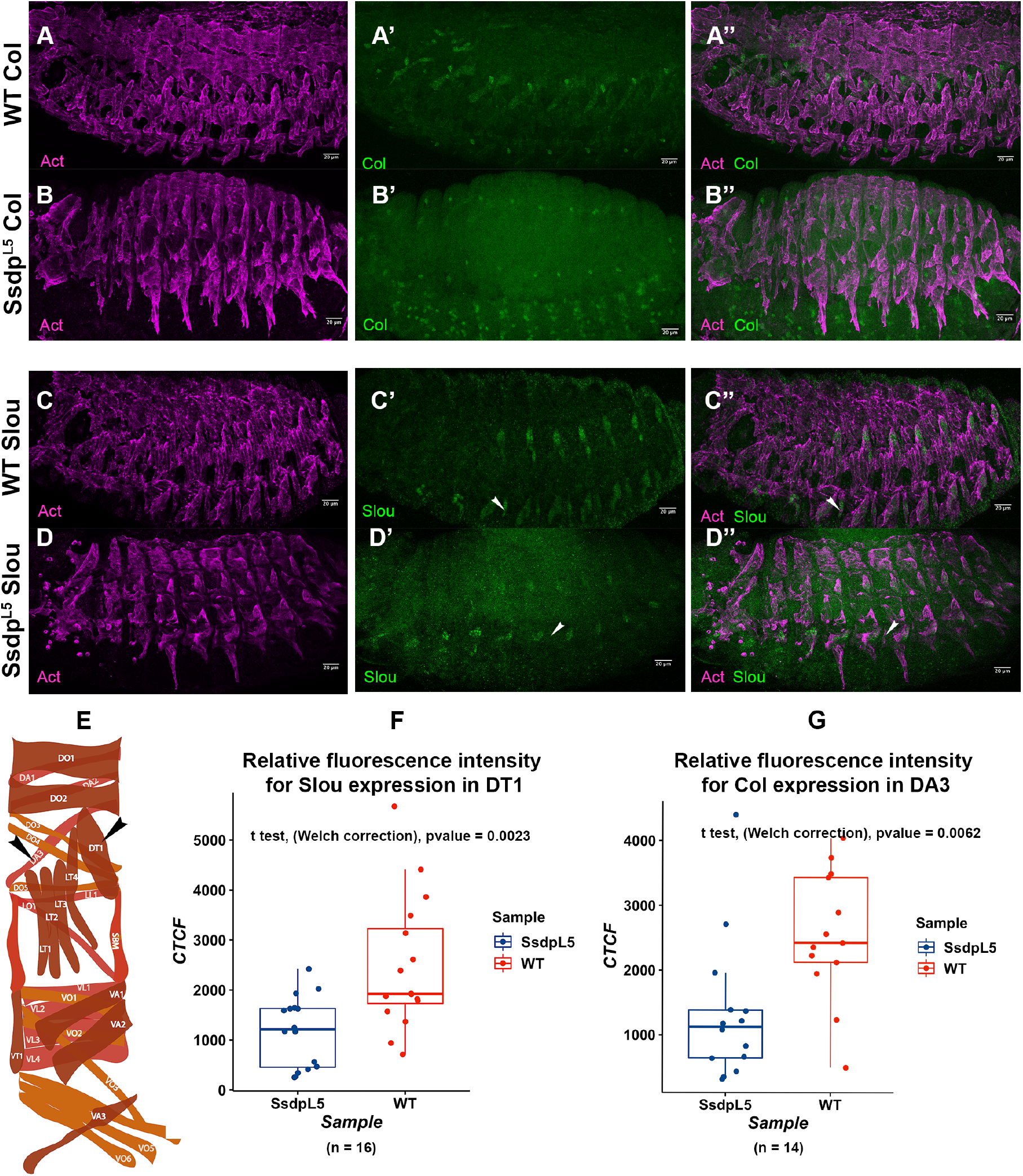
Muscle identity transcription factor (iTF) expression is downregulated in *Ssdp* mutants. **(A-A”)** Col expression as visualized by staining using an anti-Col antibody in the DA3 muscle in WT stage 16 embryos. **(B-B”)** Col expression is downregulated in *Ssdp* mutant stage 16 embryos. **(C-C”)** Slou expression as visualized by immunostaining using an anti-Slou antibody in stage 16 WT embryos. **(D-D”)** Slou expression is downregulated in *Ssdp* mutants. Expression is almost negligible in the VT1 muscles (arrowheads in D’, D”) compared to the WT muscles (arrowheads in (C’, C”). **(E)** The WT embryonic somatic muscle pattern. The DA3 and DT1 muscles are highlighted by arrowheads. **(F-G)** A quantification of intensities of Slou (F) and Col (G) by CTCF (Corrected Total Cell Fluorescence) using ImageJ shows a significant reduction in fluorescence intensities in *Ssdp* mutants with respect to WT.

The reduced levels of iTF expression in *Ssdp* mutants is significant in the context of the acquisition of muscle identity since iTFs maintain muscle-specific levels of transcription of realisator genes that are downstream of iTFs in individual muscles to generate appropriate levels of protein for each muscle to attain its specific identity (Bataillé et al. 2017).

### 3.6. Ssdp mutant muscles differentiate, but fail to acquire their final identity

Given the downregulation of iTFs and aberrant expression of the muscle differentiation marker Tm2, we wanted to clearly distinguish between a differentiation defect versus a defect in the acquisition of muscle identity in *Ssdp* mutants. To this end, we assessed whether Ssdp is required for the acquisition of two major iTF regulated properties of individual muscles, their attachment and fusion program apart from the morphological and innervation identity phenotypes observed.

#### 3.6.1. *Ssdp* loss results in muscle-specific attachment phenotypes

ßPS-integrin localizes to the tips of LT muscles at locations where they attach to their intrasegmental attachment sites as well as to the termini of muscles that attach to intersegmental attachment sites. The LT tip-associated ßPS expression is absent in *Ssdp* mutants and its localization at the intersegmental attachment sites of the severely affected ventral muscles is much less expansive in *Ssdp* mutants compared to the WT (Figure 6A-B”).

**Figure 6.**
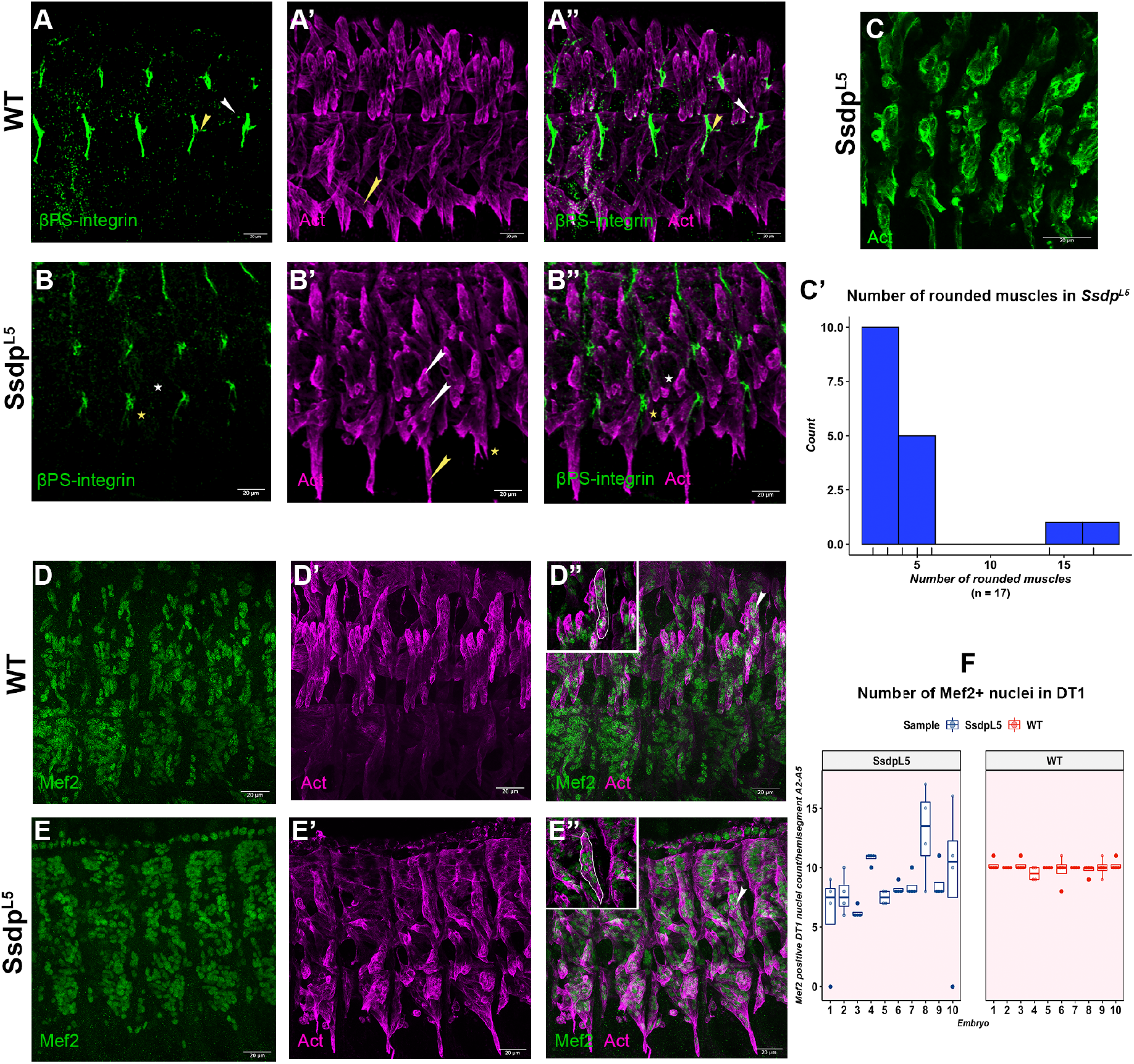
The somatic muscles of *Ssdp* mutant embryos exhibit aberrant attachment and fusion. **(A-A”)** In stage 16 WT embryos, ßPS-integrin localizes to the tips of LT muscles where they attach to intrasegmental attachment sites (white arrowhead in A, A”) and at the intersegmental attachment sites of ventral muscles (yellow arrowhead in A, A”). **(B-B”)** *Ssdp* mutants lack ßPS-integrin localization at the tips of LTs (white asterisk in B, B”) and there is much less accumulation at ventral muscle attachment sites (yellow asterisk in B, B”). Muscles that attach to intrasegmental sites such as the LTs and VA2 muscles are frequently reduced to globs (white arrowheads in B’). Unlike WT embryos where the ventral VO4-VO6 muscles traverse into the next hemisegment for attachment (arrowhead in A’), they navigate down ventrally (yellow arrowhead in B’) or fail to extend in search of their attachment sites (asterisk in B’) in *Ssdp* mutants. **(C)** In *Ssdp* mutants, around 15% of the embryos (3 out of 20) present with almost all muscles being rounded. **(C’)** Among embryos where only a portion of the muscles are rounded, the majority of embryos have 2-4 rounded muscles in hemisegments A2-A5. **(D-D”)** In the DT1 of WT stage 16 embryos, Mef2+ nuclei are localized mostly towards muscle ends (arrowhead and inset in D”). **(E-E”)** Mef2+ nuclei are clearly discernible in *Ssdp* mutants. In DT1, however, they are localized centrally (arrowhead and inset in E”). **(F)** The number of Mef2+ nuclei in the DT1 muscle in hemisegments A2-A5 displays a high degree of variation in *Ssdp* mutants, between embryos as well as within the same embryo. WT DT1 muscles have 10-11 nuclei with very little variation.

A lack of muscle extension to reach their attachment sites and misdirected ventrally extending VO4-6 have been observed in conditions where the ventral muscle iTF *vestigial* (*vg*) and its interacting partner *scalloped* (*sd*) are ectopically expressed in all somatic muscles driven by Mef2-Gal4 (Deng et al. 2009). This phenotype is also observed in *stripe* (*sr*) mutants and on ectopic expression of the *sr-b* isoform of *sr* in the ventral midline (Frommer et al. 1996; Vorbrüggen et Jäckle 1997). The ventral most VO4-6 muscles are severely affected in conditions of *Ssdp* loss of function and appear fused and indistinguishable from each other. In WT embryos, they traverse into the adjacent segment to find their attachment sites and attach to them. In *Ssdp* mutants, they either remain in the same segment and travel straight ventrally instead or do not extend at all (Figure 6A’, B’). Thus, the loss of Ssdp and the ectopic expression of Vg result in redundant ventral muscle phenotypes indicating a Ssdp-induced imbalance in yet another iTF, Vg and/or deregulated ectodermal Sr expression.

Muscles assuming a rounded appearance are observed in *sr* (Frommer et al. 1996) mutants that fail to attach and *myospheroid (mys* coding for ßPS-integrin) mutants that detach after initial attachment (Wright 1960; Leptin et al. 1989). The rounded muscle phenotype is frequently seen in *Ssdp* mutants. In 15% of embryos, almost all muscles present a rounded appearance with the lateral region being the most affected. In embryos where only a small proportion of muscles are rounded, up to 4 external muscles are rounded per embryo in hemisegments A2-A5 (Figure 6C, C’).

#### 3.6.2. Myoblast fusion is defective under the loss of *Ssdp*

Myocyte enhancer factor 2 (Mef2), the *Drosophila* homolog of vertebrate MEF2, is a MADS box transcription factor in the absence of which muscle FCs fail to differentiate after they are correctly specified (Ranganayakulu et al. 1995). It regulates the expression of a vast array of genes (Junion et al. 2005; Sandmann et al. 2006) in a dose dependent manner (Elgar, Han, et Taylor 2008). It regulates muscle identity in concert with iTFs such as Ladybird (Lb) and Vestigial (Vg) (Junion et al. 2007; Deng et al. 2009) by differentially regulating the levels of muscle genes based on its interactors. Muscle size is determined by the number of rounds of fusion, that is in turn dictated by the iTF code that regulates the muscle-specific expression levels of identity realisators including cytoskeletal modulator genes such as *Muscle Protein 20* (*Mp20*) and *Paxillin* (*Pax*) to regulate fusion (Bataillé et al. 2010). These genes start to get expressed from stage 13, which coincides with the time when we first observe detectable expression of the long isoforms of *Ssdp* in muscles.

We observe Mef2 expression in the nuclei of developing myotubes in *Ssdp* mutants similar to WT myonuclei (Figures 6D-E”). However, once the differentiation program is correctly initiated by the fusion of FCs with FCMs in *Ssdp* mutants, there are aberrations in the execution of muscle identity dependent fusion programs. In the WT, the Slou+ DT1 muscle, for example, has 10-11 myonuclei after fusion in the A2-A5 abdominal hemisegments with very little variation among embryos. *Ssdp* mutant DT1 muscles exhibit huge variations in the number of myonuclei among different embryos as well as within the same embryo ranging from a missing DT1 muscle to presenting up to 17 myonuclei (Figure 6F). In addition, DT1 presents an aberrant, elongated morphology with centralized nuclei as opposed to nuclei that localize more towards muscle ends in WT DT1 muscles (Figure 6D”, E”).

These morphology, innervation, attachment and fusion defects observed along with the attenuated, but correctly patterned expression of iTFs such as Slou and Col in expected muscles in parallel with the unaffected expression of the key muscle differentiation factor Mef2 support the view that the muscle identity program is initiated on differentiation, but the maintenance and establishment of a muscle-specific identity program is hindered in embryos lacking *Ssdp*.

### 3.7. The loss of Ssdp affects Wg expression

It was recently shown that Ssdp is involved in the transduction of Wnt/Wg signaling as part of Wnt enhanceosome complexes (Fiedler et al. 2015; Renko et al. 2019), although no function has been attributed to it during embryonic development yet. We thus asked whether Ssdp-Wg interactions could at least in part explain the complex muscle phenotypes of *Ssdp* mutants. We first tested whether embryonic Wg expression is maintained in a *Ssdp* loss of function context. In WT embryos, Wg displays a stage-specific, patterned and segmental epidermal expression (Ohlmeyer et Kalderon 1997). In *Ssdp* mutants, epidermal Wg expression is reduced during mid stages of development and is undetectable in late stage embryos (Figure 7A-D”).

**Figure 7.**
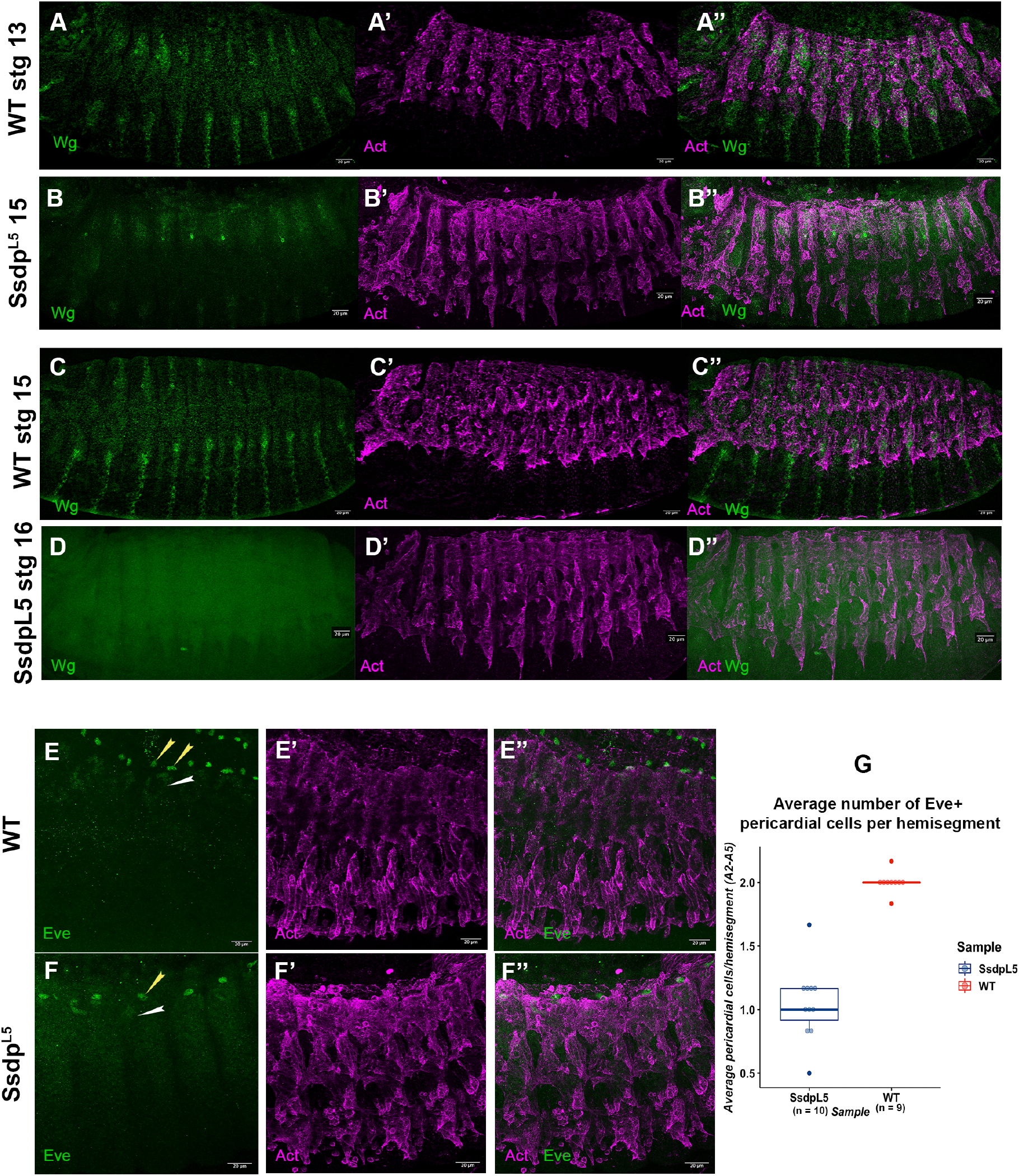
The loss of function of *Ssdp* influences Wg expression and the number of Eve+ pericardial cells. **(A-A”)** In stage 13 WT embryos, a staining with an anti-Wg antibody reveals patterned dorsal and ventral expression. **(B-B”)** In *Ssdp* mutants, Wg expression is highly reduced during mid embryonic stages where a high number of fusion foci are evident. **(C-C”)** Wg expression is maintained in stage 15 WT embryos. Its expression is more expansive ventrally. **(D-D”)** Wg expression is undetectable at later stages in *Ssdp* mutants. **(E-E”)** In WT embryos, the Eve protein is expressed in two pericardial cells per hemisegment (yellow arrowheads in E) and in the DA1 muscle (white arrowhead in E) by stage 13. **(F-F”)** In *Ssdp* mutants, we observe a single pericardial cell per hemisegment (yellow arrowhead in F). Eve expression in the DA1 muscle is highly reduced (white arrowhead in F). **(G)** A quantification of Eve+ pericardial cells in *Ssdp* mutants confirms the pericardial cell phenotype.

### 3.8. The loss of Wg and Ssdp impact Eve expression similarly

Two studies showed that the muscle and heart iTF Even-skipped (Eve) is responsive to Wg signaling. Reducing Wg signaling by expressing a dominant negative form of dTCF in the mesoderm or mutating dTCF sites in a mesodermal specific *eve* enhancer resulted in one Eve+ pericardial cell instead of the normal two per hemisegment by stage 13 (Halfon et al. 2000; Stefan Knirr et Frasch 2001). Interestingly, in *Ssdp* mutants, we observe an average of one Eve+ pericardial cell per hemisegment similar to these studies (Figure 7E-G).

These observations suggest that Ssdp is required either: i) for the maintenance of late Wg expression in the epidermis (in line with our observation of reduced epidermal Wg in *Ssdp* mutants and the epidermal expression of *Ssdp* mRNA), ii) for the transduction of Wg signals to the mesoderm as a component of the Wg enhanceosome (in line with Ssdp expression in the developing muscles) and/or iii) for both these functions.

### 3.9. Ssdp regulates a subset of Wg driven muscle characteristics and Wg plays stage-specific roles during embryonic myogenesis

In light of the similar impact of reduced *wg* and loss of *Ssdp* on the Eve iTF that is also expressed in the cardiac mesoderm, we sought to verify whether muscle defects induced by the loss of *wg* or its effector dTCF (or Pangolin (Pan)) are reminiscent of those in *Ssdp* mutants. It has previously been shown that Wg signaling mediated by dTCF is required for the proper specification of ventral muscle progenitors that give rise to the VA1, VA2 and VA3 muscles (Cox et Baylies 2005; Cox, Beckett, et Baylies 2005). Intriguingly, these muscles are severely affected in *Ssdp* mutants. VA1 and VA2 are malformed and VA3 is missing in *Ssdp* mutants (Figure 8A-B). Driving a dominant negative form of dTCF (dTCF^DN^) in all muscles using the early mesodermal 24B-Gal4 leads to partial phenotypes of missing VA1, VA2 and VA3 indicating that dTCF^DN^ is not fully penetrant in embryonic muscles. This leads to a heterogeneity in phenotypes. However, some of the muscle phenotypes including an aberrant, elongated shape or loss of DT1 as well as missing VA3 muscles are common to dTCF^DN^ and *Ssdp* mutant embryos (Figure 8B-D). Therefore, dTCF^DN^ mutant phenotypes only partially overlap that of *Ssdp* mutants.

**Figure 8.**
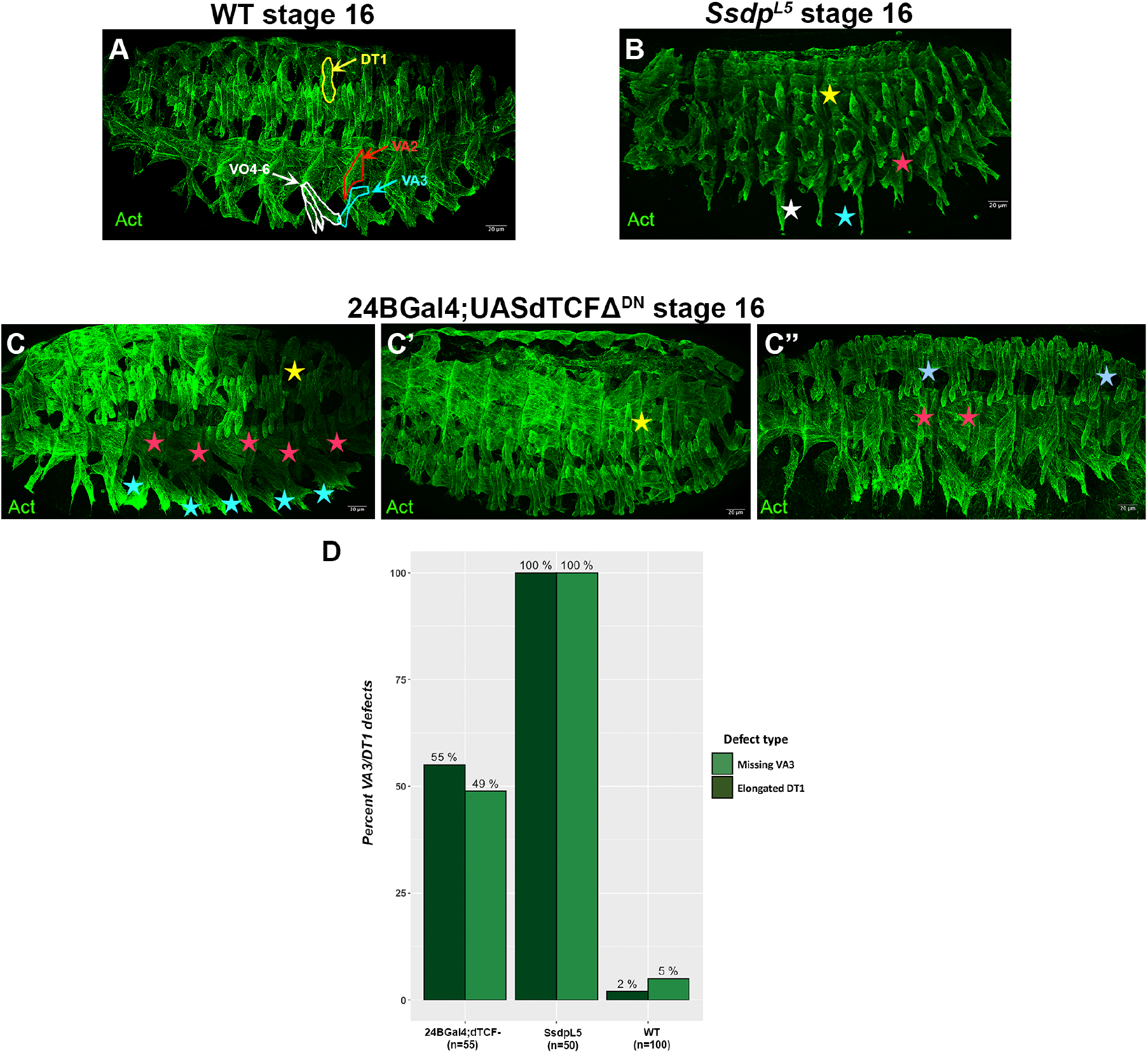
*Ssdp* mutant phenotypes are more pronounced than dTCF dominant negative mutants. **(A)** The somatic muscle pattern in stage 16 WT embryos. The DT1, VA2, VA3 and VO4-6 muscles are highlighted in yellow, red, white and cyan respectively. **(B)** The somatic muscle pattern in stage 16 *Ssdp* mutant embryos in which all of these muscles exhibit morphological defects (highlighted by similar colored asterisks as muscles in (A)). **(C)** Expressing a dominant negative form of dTCF (dTCF^DN^) using an early muscle specific driver, 24B-Gal4 reveals a heterogeneity in phenotypes depending on the penetrance of dTCF^DN^. The ventral VA1, VA2 and VA3 muscles are specified in a Wg dependent fashion and these muscles are missing in all hemisegments in C (red and cyan asterisks). In this embryo, the Slou+ DT1 muscles are missing or malformed (yellow asterisk). **(C”)** Highlighting the heterogeneity of phenotypes, this embryo has missing VA1 and/or VA2 (red asterisks) in a few hemisegments associated with LT duplications (light blue asterisks). **(C”)** In some embryos, DT1 presents an abnormal, elongated morphology (yellow asterisk) or is absent similar to *Ssdp* mutants (B). **(D)** A quantification of the missing VA3 and elongated DT1 phenotypes in dTCF^DN^ versus *Ssdp* mutants reveals that these phenotypes are present in close to 50% of hemisegments in the partially penetrant dTCF^DN^ mutants.

We also analyzed *wg* temperature sensitive mutants (*wg^ts^*) by inhibiting *wg* at different developmental stages to test Wg requirements in developing somatic muscles. When flies were allowed to develop normally at 18°C until around stage 9-11 and the eggs were then shifted to a non-permissive temperature of 28°C to inhibit *wg* expression until stage 16, we observe 2 distinct phenotypes that might correspond to different stages when *wg* was switched off in the embryo (Figure 9B, B’). The first phenotype is a complete disruption of muscle development. This is expected since Wg is required for mesoderm specification in the embryo (Azpiazu et al. 1996). The second phenotype is highly deregulated somatic muscle development with the ventral VO4-6 muscles being directed straight down ventrally instead of traversing into the adjacent hemisegment similar to *Ssdp* mutants. When *wg* is switched off between stage 15 and 16, we observe an additional phenotype where most muscles are specified correctly, but VA3 is undetectable as in the case of *Ssdp* mutants, which raises the question of whether Wg is necessary for the maintenance of this muscle (Figure 9B”). Muscles display severe attachment defects with the LT muscles extending too far dorsally to attach and ventral muscles associating with incorrect attachment sites. VA1 and VA2 present aberrant morphologies that could be the result of them attaching to ectopic sites. Thus, Wg plays stage-specific roles during embryonic somatic muscle development.

**Figure 9.**
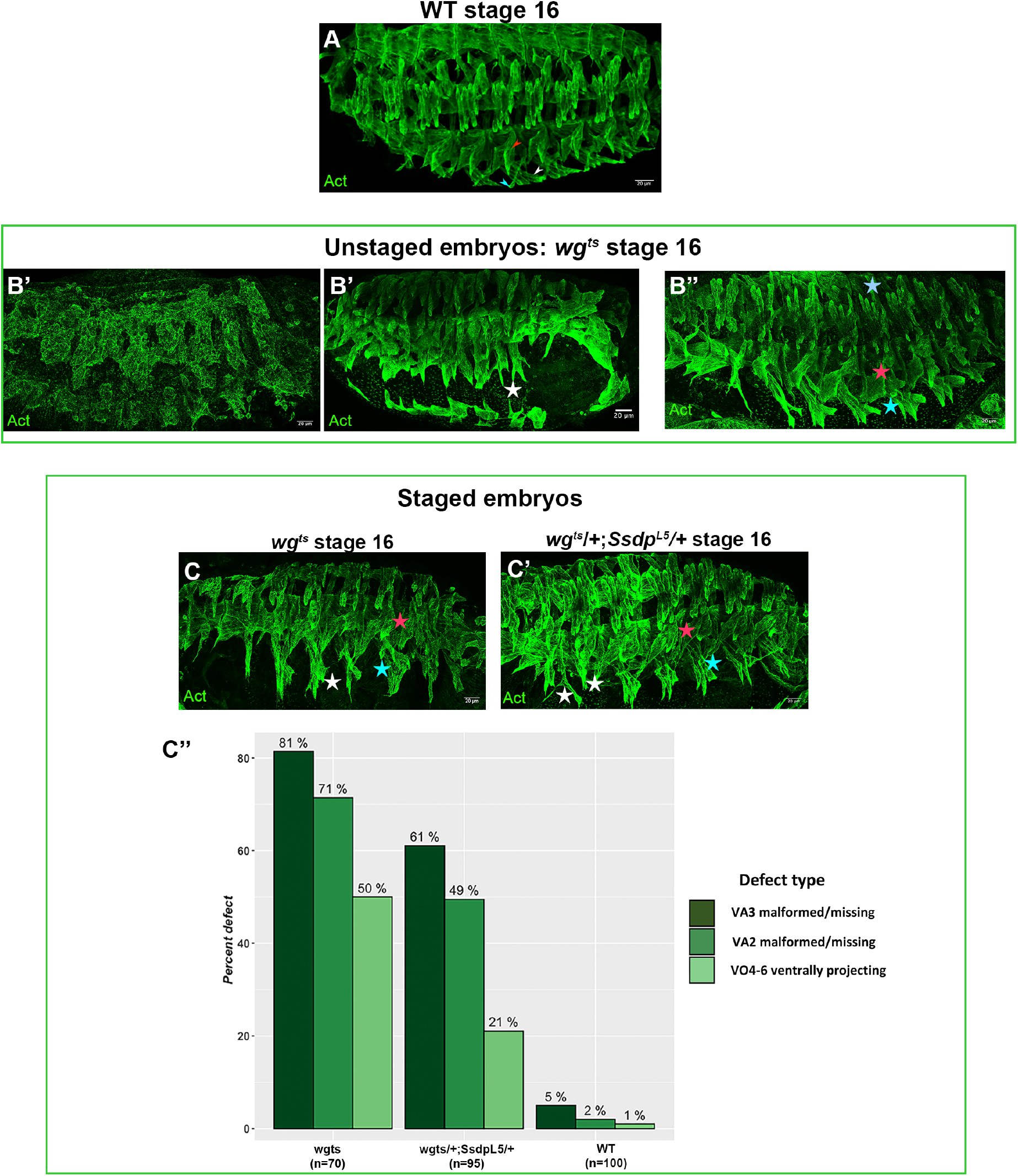
Wg plays stage-specific roles during embryonic somatic muscle development and genetically interacts with Ssdp during mid stages to regulate VA2 and VA3 identity. **(A)** The somatic muscle pattern in stage 16 WT embryos. VA2, VA3 and VO4 are highlighted (red, cyan and white arrowheads respectively). **(B-B”)** Unstaged *wg^ts^* mutant embryos with 0-12 hours of normal development with continuous egg-laying at 18°C (~stage 9-11) before shifting to the non-permissive 28°C to deactivate *wg* until stage 16 are displayed in (B) and (B’). They display multiple phenotypes including completely disrupted myogenesis (B) and deregulated myogenesis with ventrally projecting VO4-6 (B’) also observed in *Ssdp* mutants. When allowed to develop normally for longer (0-26 hours or ~stage 15-16) before being shifted to 28°C (B”), additional severe attachment phenotypes are observed in VA2 and LTs (red and light blue asterisks in B” respectively). VA3 is undetectable (cyan asterisk). **(C)** Staged *wg^ts^* mutant embryos with 3 hours of egg laying at 18°C followed by normal development for 13-17 hours (~stage 12-13) before shifting to 28°C exhibit specific phenotypes. VA2 and VA3 are malformed/missing while VO4-6 are indistinguishable from each other and project ventrally similar to *Ssdp* mutants. **(C’)** Similarly staged *wg^ts^-/+;Ssdp^L5^-/+* transheterozygotes display a disorganized actin cytoskeleton. They display attachment defects in the ventral-muscles in anterior hemisegments (white asterisks). Some phenotypes such as missing or malformed VA2 and VA3 muscles (red and cyan asterisks respectively) overlap those observed in *wg^ts^* mutants. **(C”)** A quantification of ventral muscle defects in staged *wg^ts^* mutants and *wg^ts^/+;Ssdp^L5^/+* transheterozygotes reveals that the percentages of these defects are significantly higher compared to the WT.

We analyzed precisely staged *wg^ts^* embryos to verify if there were specific phenotypes that were distinguishable. When flies were allowed to lay eggs for 3 hours at the permissive temperature and the eggs were allowed to develop normally until around stage 12-13, which is right after FC specification, before being shifted to the non-permissive temperature of 28°C, we observe distinct phenotypes (Figure 9C). LT muscles extend to much greater distances than WT embryos. The VO4-6 muscles project ventrally in search of attachment sites as observed in *Ssdp* mutants in many cases. The VA2 and VA3 muscles are smaller and present aberrant morphologies. Given the temporal specificity of phenotypes observed, we examined similarly staged *wg^ts^*/+;*Ssdp^L5^*/+ transheterozygotes to determine possible genetic interactions between Ssdp and Wg (Figure 9C’). These embryos display heterogenous actin staining indicating a disorganized actin cytoskeleton as seen in *Ssdp* mutants. They exhibit severe morphological and attachment defects in VO4-6 in the anterior-most hemisegments. Somatic muscles appear relatively normal in some embryos, but ventral muscles exhibit defects in morphology and attachment. VA2 and VA3 are absent or smaller with aberrant morphology also observed in similarly staged *wg^ts^* mutants. A quantification of these phenotypes in staged embryos (Figure 9C”) reveals that ventral defects are more pronounced in *wg^ts^* mutants compared to *wg^ts^*/+;*Ssdp^L5^*/+ transheterozygotes with VA3 being the most affected, followed by VA2.

These observations suggest that all *Ssdp* loss of function phenotypes cannot be explained by muscular transduction of Wg signals by dTCF. However, one has to take into consideration that dTCF^DN^ is not fully penetrant in embryonic muscles. *wg^ts^* mutants, on the other hand, present more severe phenotypes. There is distinct overlap between phenotypes observed in *Ssdp* mutants versus dTCF^DN^ mutants and *wg^ts^* mutants. The analysis of staged *wg^ts^*/+;*Ssdp^L5^*/+ transheterozygotes indicates a genetic interaction between Wg and Ssdp at mid developmental stages to regulate actin cytoskeletal dynamics and the establishment of muscle identity of a specific subset of ventral muscles given that heterozygotes for each of them do not display these phenotypes. These observations suggest a stage dependent requirement and specific interaction between Wg signaling and Ssdp in the regulation of the identity of the VA2, VA3 and VO4-6 muscles during embryonic somatic muscle development.

## 4. Discussion

### Ssdp and isoform specificity during somatic myogenesis

In line with the differential expression of *Ssdp* in muscles in our transcriptomic data, our analysis of *Ssdp* mutants shows that this gene plays a critical role during muscle development. The differential expression of *Ssdp* mRNA under translation among different muscle subsets in our transcriptomic data reveals a potential muscle-subset-specific role for this gene. This might be due to its requirement to form specific complexes with LIM homeodomain factors such as the LT iTF, Ap (Bronstein et al. 2010). The observation of an enrichment for CT-rich motifs in the LT subset is also in line with the significant upregulation of *Ssdp* under translation in our dataset. No direct DNA-binding has been proven *in vivo* for Ssdp. It has been shown to bind DNA by interacting with its cofactors as part of the ChiLS and Wnt enhanceosome complexes. So, it is possible that this enrichment represents Ssdp’s interaction with other cofactors. Trl (Trithorax-like) is a potential interactor given that it is known to bind complementary GA-rich motifs and is significantly upregulated along with Ssdp during T3 in our data.

In humans, *SSBP3* downregulation and mis-splicing with the retention of exon 6 has been observed in the muscles of myotonic dystrophy type 1 and 2 patients as well as patients with neuromuscular diseases (NMD) such as Duchenne muscular dystrophy (Bachinski et al. 2014), although it has received no attention and no role during myogenesis has been attributed to it yet. The differences in the expression patterns of the *in situ* probes targeting the long isoforms versus all isoforms indicate the presence of predominant isoforms with a potential isoform switch at different stages of development, which could be related to isoform specific functionality already hinted at in human muscular dystrophies.

### Ssdp and its influence on muscle identity properties

Identified methods by which iTFs direct identity acquisition include controlling the muscle-specific number of rounds of myoblast fusion (Bataillé et al. 2010), reprogramming newly fused nuclei to adopt a muscle-specific program (Bataillé et al. 2017) and correct attachment site selection (Carayon et al. 2020). Given this, our observations of downregulated iTF expression as well as deregulated myoblast fusion and attachment in specific muscles suggest a requirement for Ssdp during mid myogenesis after the muscle identity program and muscle differentiation have been initiated. This is when we detect expression of the long *Ssdp* isoforms in somatic muscles, which is potentially related to muscle and stage-specific functions for these isoforms. However, this does not rule out a requirement for Ssdp during early stages of development when it is ubiquitously expressed and the zygotic null mutants used here might hamper the observation of this early requirement.

### The roles of Ssdp and Wg on muscle phenotypes

The extremely low Wg expression in *Ssdp* mutants during mid stages and lack of Wg observed at later stages indicate a role for Ssdp in the maintenance of Wg expression at these stages when the long isoforms are highly expressed. Since the Wg dependent Slou+ cluster is largely specified except for VA3, this indicates sufficient Wg levels at early stages of muscle development when VA1/2 progenitors are specified, but a dependence on Ssdp for VA3 specification. It is unclear if the low levels of iTF expression observed at later stages is the consequence of downregulated Wg. Given that Ssdp is part of the evolutionarily conserved canonical Wnt enhanceosome, it is not surprising that it in turn affects Wg expression as this study unveils for the first time. Since the vertebrate Wnt genes play very specific roles during myogenesis, it remains to be seen if Ssdp also influences the expression of other *Drosophila* Wnts and if they have roles during myogenesis. The analysis of *wg^ts^*/+;*Ssdp^L5^*/+ transheterozygotes confirms stage and muscle specific interactions of Ssdp to regulate Wg signaling, especially in establishing the identity of VA2 and VA3. Whether this is via the canonical pathway is an open question.

This study also reveals stage-specific roles for Wg during all stages of embryonic somatic myogenesis for the first time. Although Wg is known to be required for the specification of the mesoderm and the specification of some FCs during early stages, no role has been attributed to it during mid-late stages. We identify a role for it in regulating the correct attachment of muscles during later stages as evidenced by the severe attachment phenotypes observed when *wg* is deactivated during late stages.

### Singular or converging roles of Ssdp could play a part in somatic muscle development

As a component of the Wnt enhanceosome and because Wnt signaling is a central component of the developmental symphony, any deregulations in components of this pathway can have deleterious cascading effects. Since humans have 3 *SSBP* family members and *Drosophila* only one, it could mean that the single gene carries out all functions associated with vertebrate counterparts or that *Drosophila* requires only a subset of its vertebrate counterpart family’s functionality.

The stark variability in the number of myonuclei in the DT1 muscle suggests a stochasticity due to deregulations in protein stoichiometry. So, Ssdp might play a role in providing a context-dependent boost of gene transcription. Given the incomplete overlap of phenotypes in various contexts in our study, it could be the known role of Ssdp as part of ChiLS that triggers the defects observed or a combination and convergence of this with unknown Ssdp roles that have a cumulative effect on myogenesis. Whether Ssdp plays cytoplasmic roles apart from forming nuclear complexes is unknown. The LisH motif (van Meyel, Thomas, et Agulnick 2003) present in Ssdp has been implicated in controlling MT dynamics by homodimerizing or dimerizing in trans with other LisH proteins as well as in regulating protein localization (Emes et Ponting 2001; M. H. Kim et al. 2004; Gerlitz et al. 2005; Kannan et al. 2017). Analyzing Ssdp cofactors and conserved regions in the Ssdp protein other than the LUFS domain would help elaborate on Ssdp functions.

## Author Contributions

Study conceptualization, P.P. and K.J.; methodology, P.P. and K.J.; software, P.P.; validation, P.P.; formal analysis, P.P.; investigation, P.P.; resources, P.P. and K.J. with help from J.P.; data curation, P.P. and K.J.; writing–original draft preparation, P.P.; writing–review and editing, P.P. and K.J.; supervision, K.J.; project administration, K.J.; funding acquisition, K.J. All authors have read and agreed to the published version of the manuscript.

## Acknowledgments

We would like to thank Benjamin Bertin for generating the TRAP datasets used for bioinformatic analysis in this study.

