## Supplementary data for "Ssdp influences Wg expression and embryonic somatic muscle identity in *Drosophila melanogaster*"

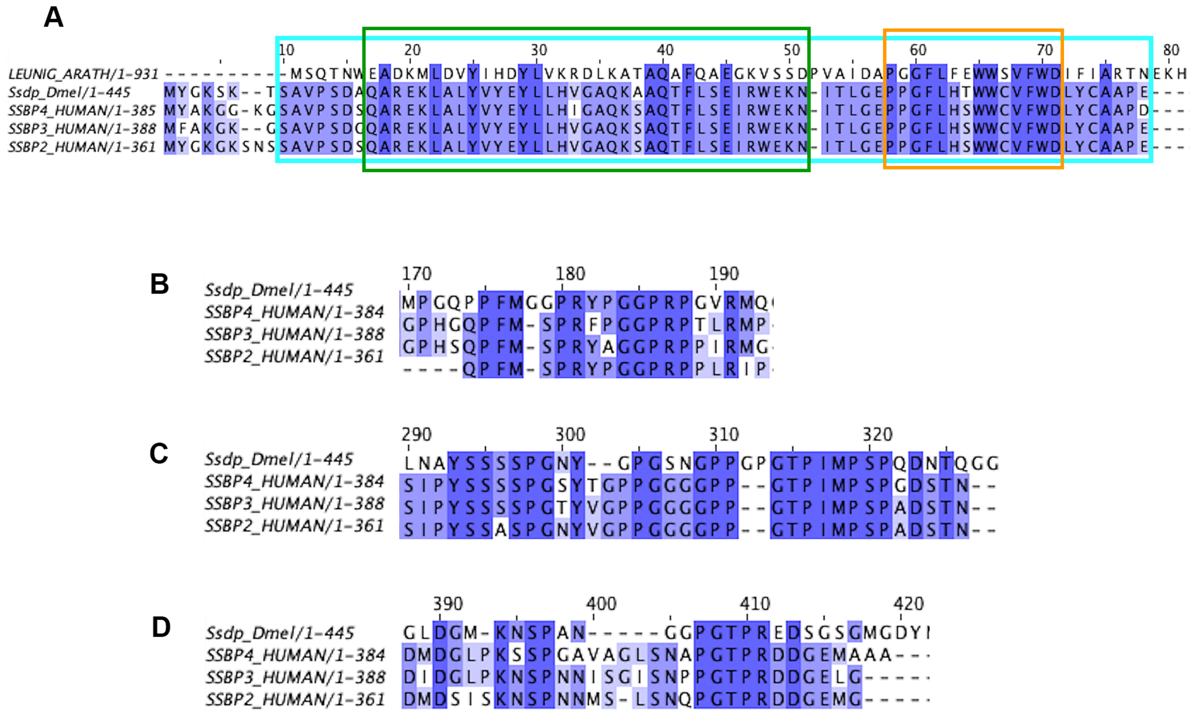

**Supplementary figure 1.**

**Ssdp has four regions with highly conserved amino acid residues when aligned against human SSBPs.**

**(A)** The first 80 residues of the Ssdp protein aligned with the conserved LUFS domain of the LEUNIG protein in *Arabidopsis thaliana* and canonical isoforms of the human homologues, SSBP2, SSBP3 and SSBP4. The LUFS domain that is highlighted with a cyan rectangle permits the formation of a complex with Ssdp partner, Chi (LDB1 in humans). Within it is a LisH domain (green rectangle) and another conserved domain (orange rectangle) of unknown function. **(B-D)** Apart from the already identified LUFS domain, the alignment unveils three other regions with blocks of highly conserved amino acid residues. (B) is part of a proline-rich stretch that was identified as being essential for fore and midbrain development in mice.

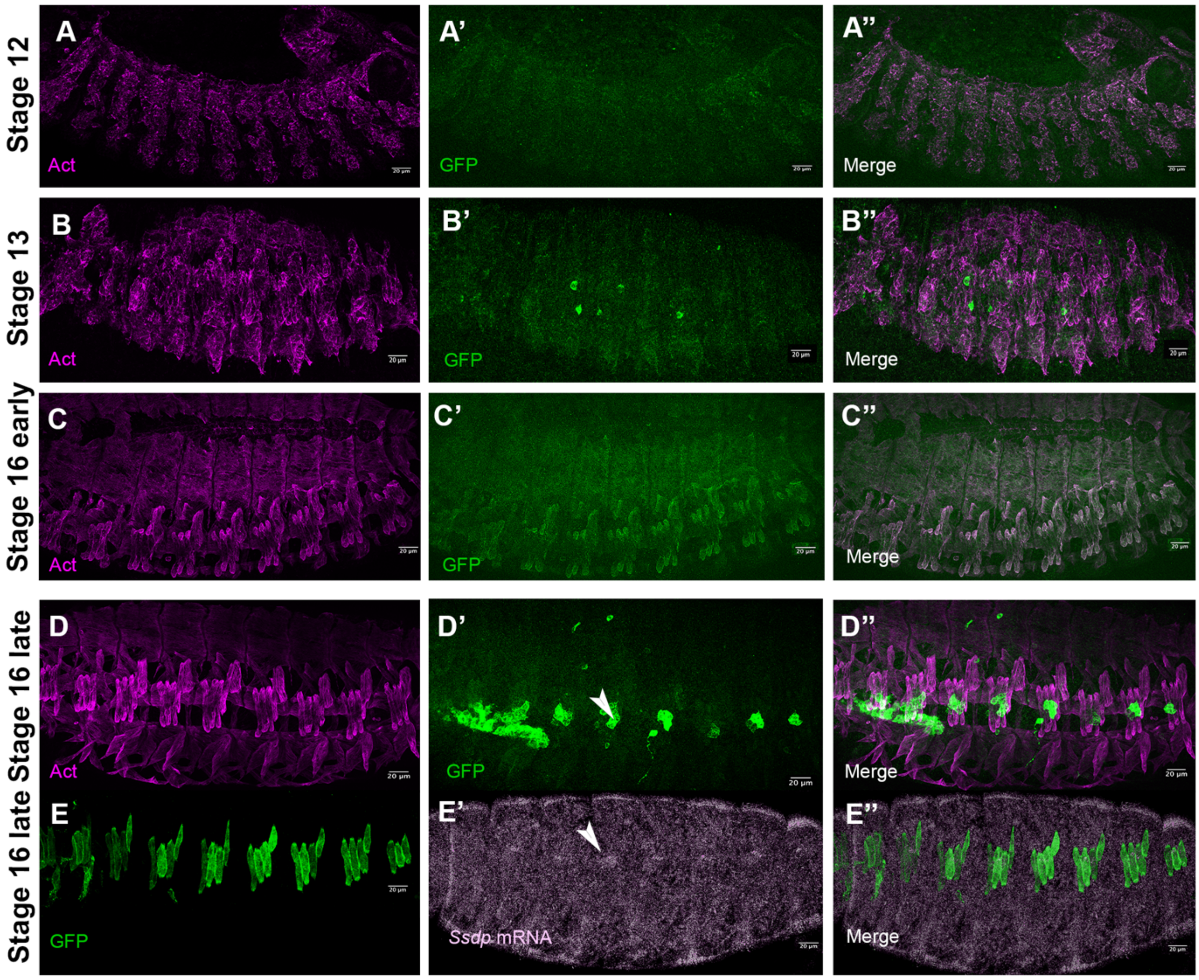


**Supplementary figure 2.**

**Ssdp protein expression as revealed by a Ssdp-Gal4 driven UASAct5CGFP.**

**(A-A'')** Low levels of GFP can be detected in the dorsal muscles by stage 12. **(B-B'')** All muscles express GFP by stage 13. **(C-C'')** Expression is much stronger at early stage 16 with marked GFP expression in the Lms+ LT and Slou+ DT1 muscles. **(D-D'')** By late stage 16, a high GFP signal is detected in the sub-epidermal chordotonal organ (arrowhead) that acts as a proprioceptor and is situated just above the LT muscles. **(E-E'')** RNA FISH of *Ssdp* transcripts reveals similar high expression levels of mRNA in this region (arrowhead) above the LT muscles.
